# How is structural divergence related to evolutionary information?

**DOI:** 10.1101/196782

**Authors:** Diego Javier Zea, Alexander Miguel Monzon, Gustavo Parisi, Cristina Marino-Buslje

## Abstract

Conservation and covariation measures, as other evolutionary analysis, require a high number of distant homologous sequences, therefore a lot of structural divergence can be expected in such divergent alignments. However, most works linking evolutionary and structural information use a single structure ignoring the structural variability inside a protein family. That common practice seems unrealistic to the light of this work.

In this work we studied how structural divergence affects conservation and covariation estimations. We uncover that, within a protein family, ~51% of multiple sequence alignment columns change their exposed/buried status between structures. Also, ~53% of residue pairs that are in contact in one structure are not in contact in another structure from the same family. We found out that residue conservation is not directly related to the relative solvent accessible surface area of a single protein structure. Using information from all the available structures rather than from a single representative structure gives more confidence in the structural interpretation of the evolutionary signals. That is particularly important for diverse multiple sequence alignments, where structures can drastically differ. High covariation scores tend to indicate residue contacts that are conserved in the family, therefore, are not suitable to find protein/conformer specific contacts.

Our results suggest that structural divergence should be considered for a better understanding of protein function, to transfer annotation by homology and to model protein evolution.

## 1. Introduction

Since the experiments of Anfinsen, we have known that protein structural information is codified in a protein sequence (Anfinsen, 1973). However, predicting which positions determine structural arrangements and functional features remains difficult when using information contained in a single sequence. A convenient way to solve this problem is to use well populated Multiple Sequence Alignments (MSAs) of homologous proteins to derive evolutionary information. Specifically, highly conserved or co-evolving positions can give us an idea of functionally and/or structurally important protein sites.

Conserved positions in an MSA are thought to evolve under several constraints as described by the neutral theory of molecular evolution (Kimura, 1983). According to this theory, most evolutionary changes result from mutations with minimal or minor functional impact that are fixed via random genetic drift. However, mutations in biologically important residues (e.g., active sites) that impair protein function, folding or stability, are either subsequently eliminated by purifying selection or compensated for by other mutations elsewhere in the protein. Within protein and genetic studies, the concept of conservation is heavily employed to infer structural and/or functional roles (Cooper and Brown, 2008; Guharoy and Chakrabarti, 2005; Karlin and Brocchieri, 1996; Lichtarge et al., 1996; Worth et al., 2009). Several methods to measure conservation are based on information theory (Valdar, 2002), such as Shannon entropy (EN) and Kullback–Leibler (KL) divergence (Johansson and Toh, 2010).

The other source of evolutionary information in a protein family’s MSA is coevolution, which basically describes the interdependence of the substitution pattern between different positions (de Juan et al., 2013). Since structural and/or functional constraints involving multiple non-independent protein sites can lead to coevolutionary signals, covariation measures are used as a proxy for coevolution. Mutual Information (MI) between positions is a covariation method,coming from information theory, and is optimised to detect coevolution (Buslje et al., 2009). Other methods try to avoid transitive covariation signals, such as, for example, Gaussian Direct Coupling Analysis (GCA) (Baldassi et al., 2014). Covariation measures can nowadays be used to infer tertiary contacts (Buslje et al., 2009; Jones et al., 2012; Morcos et al., 2011), functional sites (Marino Buslje et al., 2010) and protein–protein interactions (Hopf et al., 2014; Ovchinnikov et al., 2014).

Most works linking evolutionary and structural information use a single structure to represent the structural space of a given MSA. Some of these works analyse a particular protein of interest, while others use evolutionary information to derive general trends in proteins such as, for example, the prediction of binding sites (Liang et al., 2006), the location of catalytic sites (Capra and Singh, 2007; Marino Buslje et al., 2010), and tertiary contacts and fold prediction (Sutto et al., 2015). Other studies focus on protein evolution, trying to find functional and structural constraints (Worth et al., 2009). In particular, the Relative Solvent Accessible Surface Area (RASA)(Franzosa and Xia, 2009) and local packing density (S. W. Yeh et al., 2014) were identified as the main determinants of evolutionary rates. For all these studies, the use of a single structure to represent the whole structural space of proteins in an MSA seems unrealistic.It is well known that to gain confidence in conservation and covariation, MSAs with a high number of distant homologous sequences are required. Well populated and divergent Pfam alignments are a common choice for measuring these variables (Simonetti et al., 2013). A lot of structural divergence can be expected in such divergent MSA because the relationship between sequence identity and structural divergence is exponential (Chothia and Lesk, 1986a; Flores et al., 1993; Russell et al., 1997). Moreover, structural differences between proteins of the same (same MSA) may arise not only because of evolutionary divergence due to the accumulation of mutations, but also because of the conformational diversity of a single protein sequence. The extent of conformational diversity, the structural differences between conformers describing the native state of a protein, can be as high as 23.4 Å of RMSD (Burra et al., 2009).The observed difference between conformers may be due to secondary and tertiary element arrangements (Gerstein and Krebs, 1998) as well as loop movements (Gu et al., 2015), for example.

In this context, it is interesting to think about how expected structural differences in an MSA are reflected in sequence-based evolutionary information derived from conservation (a proxy to low evolutionary rates) and coevolution. The relative accessible surface area and contact density have received enormous attention as the main structural factors determining evolutionary rates (for a review see (Echave et al., 2016)). In general, these studies were carried out using the RASA and contact density from a single structure without taking into account the variability of these variables inside a protein family. This was even more extreme in the case of covariation scores, where inter-residue contacts were used to optimise their parameters or to compare the performance of different methods (Buslje et al., 2009; Jones et al., 2012; Zea et al., 2016), even when residue contacts may vary between protein conformers and, even more so, between different protein structures.

In this work, we studied how evolutionary information measures on Pfam alignments are related to structural information when all available structures are used. In particular, we were interested in measuring how much change occurred in the correlation of conservation against the RASA and the performance of the prediction of inter-residue contact of covariation methods when different structures from an MSA were used. We also wanted to know how structural variables are related to evolutionary variables, taking into account the RASA and inter-residue contacts of family all available structures.

## 2. Materials and methods

### 2.1. Data set construction

We performed all the analysis using ad hoc scripts in the Julia language, taking advantage of the MIToS toolkit (Zea et al., 2016). We selected Pfam protein families (version **30.0**) (Finn et al., 2016) with at least 2 proteins with high resolution structures (resolution < 3 Ångströms) and a corresponding SIFTS file (Velankar et al., 2013). We checked the identity between the PDB ATOM sequence and the uniprot sequence in the MSAs. Also structures should cover at least 80% of the MSA aligned columns, and be longer than 30 residues. Columns having missing residues are not used in this analysis. In such a way, we ended up with **1808** protein familiessuitable for this analysis. To ensure a good sampling of the structural space of each MSA, we cluster the sequences at 62% identity using the Hobohm I algorithm (Buslje et al., 2009;Hobohm et al., 1992) and used the subset of families with at least 4 sequence clusters with structural information (at least 1 PDB each cluster) (see supplementary material). The final dataset has **817** Pfam families. This dataset is divided into an exploration set (**245** families) and a testing or confirmatory set (**572** families) in order to avoid the post hoc theorizing (Leung, 2011). All the exploratory analysis was performed on the exploration dataset, but this work only presents the values obtained with the confirmatory subset, nevertheless, both dataset give coincident results. The confirmatory dataset was also used for the creation of each figure.Datasets, together with descriptive and measured values, are available in the supplementary files.

### 2.2. Folding coverage as CATH classification

In order to measure the coverage of the structural space of our confirmatory dataset, we counted how many classes (C) and architectures (A) of the CATH hierarchy are present. We used the SIFTS mapping at residue level to obtain the CATH id of each residue for each PDB in each MSA of the confirmatory dataset. A CATH level was assigned if only one CATH domain is present in the residues of the Pfam domain of a PDB chain. Given the differences between the Pfam domains defined at sequence level and the CATH domains defined at structure level, we could only assign a CATH domain for 68.26% of the studied Pfam domain structures. We concluded that the confirmatory dataset have a good coverage as it has members of 4/5 of the class 1 architectures (mainly alpha), 13/20 of the class 2 architectures (mainly beta), 10/14 of the class 3 architectures (alpha beta) and finally, members of the unique architecture of the class 4 (few secondary structure).

### 2.3. Structural and evolutionary analysis

For each column in the MSA we calculate the Kullback-Leibler divergence (KL) (with BLOSUM62 background frequencies) (Capra and Singh, 2007) as measure of conservation and the Shannon entropy as a measure of variation. For each pair of columns we calculate MIp (Dunn et al., 2008), ZMIp (Buslje et al., 2009) and GaussDCA (Baldassi et al., 2014) as covariation measures.

The RASA was calculated using NACCESS with default parameters as the percent of solvent accessibility of a residue in the structure compared to the solvent accessibility of that residue type in an extended ALA-x-ALA tripeptide (Hubbard and Thornton, 1993). We considered a residue as exposed if it 's RASA is greater than 20% and buried otherwise (Holbrook et al., 1990). Contacting residues were defined as those having any heavy atom at a distance <= 6.05 Ångströmg (Bickerton et al., 2011). The RMSD and the root mean square fluctuation (RMSF) values between any two structures within a protein family were calculated using the MSA as a guide for a rigid structural superimposition with the Kabsch algorithm (Kabsch, 1978, 1976). Structures were hierarchically clustered at 0.4 Ångströms (Burra et al., 2009) with the complete-linkage algorithm (Olson, 1995) to reduce redundancy and every structure was weighted as one over the number of structures in the cluster. We have used this cutoff since it is the level of the experimental error (X-ray crystallography). We calculated the probability of two residues of being in contact as the number of structures where this pair of residues are in contact, divided by the total number of structures in the MSA. For each residue, we also calculated the probability of being exposed in the MSA as the number of structures where the residue is exposed divided by the total number of structures in the MSA. For one-structure calculations, we selected one PDB from the sequence with more structural conformers of the family as reference.

### 2.4. Fraction of explained variance

Using our exploratory dataset of 245 Pfam families, we looked for linear correlations between the Kullback-Leibler divergence and the RASA of a single structure of the MSA, the weighted mean RASA of each MSA column and the weighted probability of a given position of the MSA to be exposed. The last two variables include information from all the structures in the MSA. The mean and the probability are weighted, being the weight of each structure 1 over the number of structures in a cluster, after a structural hierarchical clustering at 0.4 Å RMSD. This was done in order to avoid structural redundancy. To look for a linear correlation inside protein families between the three variable related to solvent exposure and KL we measured how much protein validates the linear model assumptions. The mean RASA and exposure probability are not linearly correlated without transformations because only 6 families validate the linear assumptions. Applying the function log(x+0.05) to transform the KL values to logarithmic scale, 63 out of 245 families validate all the model assumptions (Peña and Slate, 2006) for the three variable pairs.

## 3. Results

### 3.1. Dataset description

An MSA does not exist for which all structural conformers are known for each of its proteins. As a first step, we selected all the Pfam families that had at least two proteins with high resolution structures, ending up with 1808 families. As a second step, within each Pfam family we clustered sequences at 62% identity and counted the number sequence clusters with at least one solved structure.

We speculated that the more available structures a protein family had, the more likely we were to find greater structural differences. In order to cover the conformational space of an MSA, we selected those families with at least four sequence clusters with structural information. The minimum value of four clusters was a trade-off between retaining a sufficient number of families for the study and having enough structural information to have mean C-alpha root-mean-square deviation (RMSD) values independent of the number of sequence clusters with structures (Fig.S1. shows a boxplot of mean RMSD, depending on the number of clusters with structures).In fact, we found that the mean RMSD increased with the number of sequence clusters with structures to finally reach an asymptote.

After an statistical analysis, we considered that MSAs in our dataset should contain at least four proteins with high resolution structures belonging to four clusters at 62% sequence identity (see Supplementary information “Dependence of the mean RMSD with the number of clusters with structures”). The final dataset contains 817 protein families which we randomly split into two datasets: an exploratory one with 245 families and a confirmatory one with 572 families. All the informed results in the following sections are from the confirmatory dataset, though the results in both datasets are equivalent. This procedure allows the finding of general trends not overfitted to the available data. We also tested the coverage of CATH domains in order to ensure that our confirmatory dataset is representative of the protein fold space (see Materials and methods). MSAs of the confirmatory dataset (572 families) had 9730 sequences on average. These populated families were also diverse, having a mean identity percentage of 26.76%, on average, between sequences. The median of the mean sequence identity percentage per MSA was 25.5%, with first and third quartiles of 21.6% and 30.8%, respectively. The number of sequences and mean identity percentage distributions are shown in supplementary Fig. S2. The number of diverse sequences is the number of sequence clusters at 62% identity after a Hobohm I clustering. A corrected mutual information method performs well with 400 or more sequence clusters (Buslje et al., 2009). MSAs from our confirmatory dataset had a median of 1421 sequence clusters at 62% sequence identity, with first and third quartiles of 653 and 2913.5 sequence clusters, respectively.

### 3.2. On structural divergence, residue exposure and contacts of different structures within a protein family

Structural divergence increases as sequence identity decreases as shown in Fig. 1. This is in After agreement with the pioneering work of Chothia and Lesk, who showed that an exponential relation between structural and sequence changes (Chothia and Lesk, 1986b). In our confirmatory dataset, the Spearman rank correlation coefficient (rho) between the sequence identity percentage and the RMSD is -0.80. This correlation was calculated comparing all possible structure pairs within each protein family. Fig. 1 shows the distribution of the identity percentage and RMSD values for each structure pair belonging to the same protein family,including conformers for the same protein. The marginal distributions show that most of the structure pairs in the dataset have low sequence identity and high RMSD values. The maximum RMSD between structures is 5.65 Å, on average, and the mean RMSD is 2.48 Å, on average, calculated over all MSAs.

**Fig. 1.**
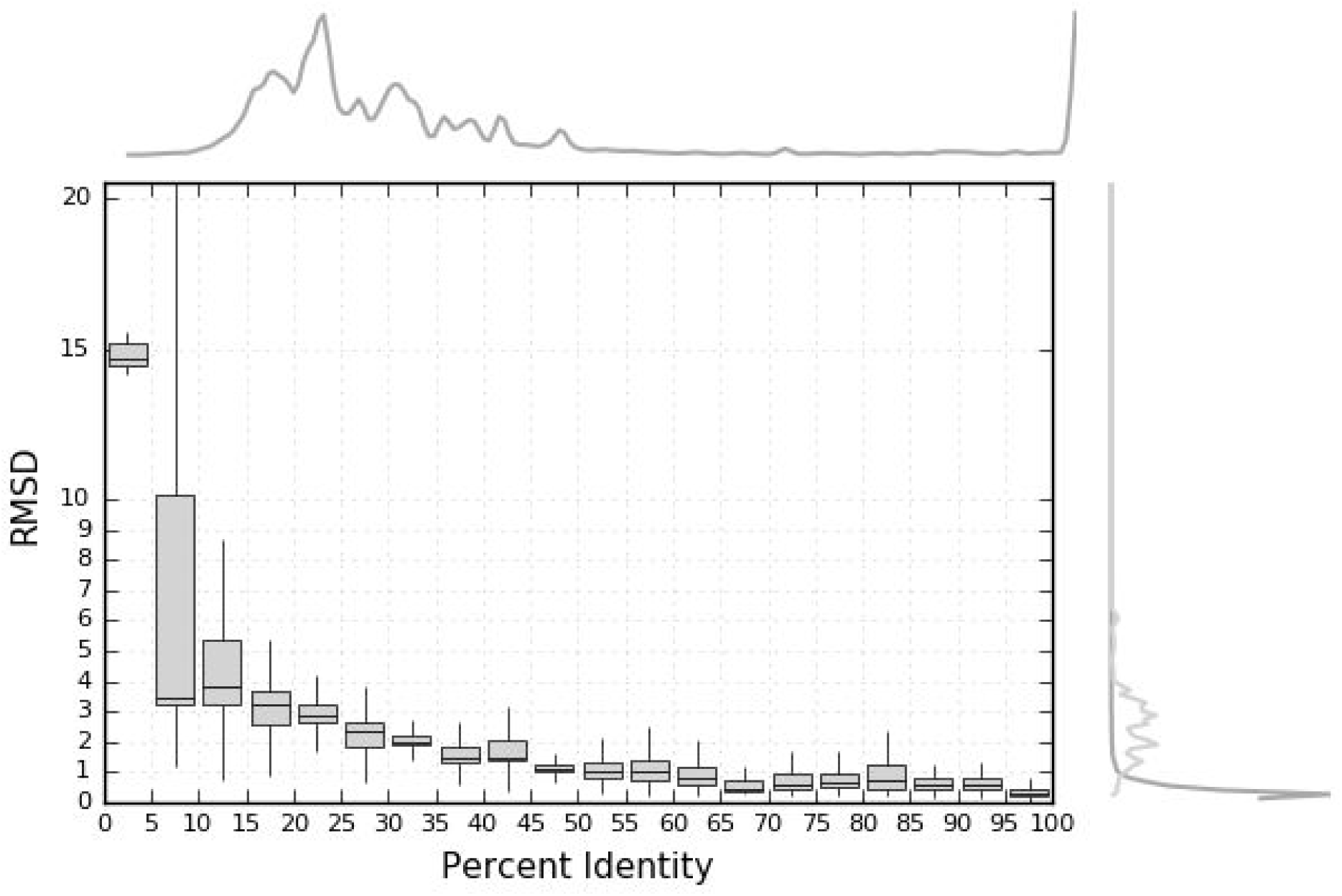
The structural divergence increases whilst the identity percent between sequences decreases. Each pair of structures inside a protein family is compared sequentially and structurally. Each box plot shows the distribution of alpha carbon RMSD at each bin of sequence identity (5% range). Outliers are not shown. RMSD is calculated after a rigid superimposition where residues (that belong to the same MSA column) are considered as structurally equivalent. The top density plot shows the distribution of sequence identity percent in the dataset. The right density plots shows RMSD distributions of protein pairs (light grey) and conformational diversity pairs (dark grey).

A reasonable expectation was that alignments with higher sequence divergence would also be more structurally divergent. The structure and sequence divergence of an MSA can be described by the mean RMSD between its structures and the mean sequence identity between its sequences, respectively (Fig. S3. shows the distribution of these variables in the confirmatory dataset). We therefore calculated the Spearman’s rho between these two variables for all the families in the dataset. We found a moderate correlation score of -0.48, which agreed with our expectation.

Within each protein family, we compared all available structures against all uncovering that the average mean change in the RASA per residue between structures was 37.39%. Also, as a consequence of the structural divergence inside a protein family, 50.63% of MSA columns, on average, changed their exposed/buried status. Furthermore, on average, 53.32% of column pairs that were in physical contact in at least one structure, were not in contact in another structure from the same MSA. These results show that the RASA and residue contacts change considerably between different structures of the same protein family.

We hypothesised that the more available structures for a protein family, the more likely we were to find greater differences between any two structures. However, since we selected those families with at least four clusters with known structure, we expected to observe a low of the RMSD with the total number of structures. In fact, Spearman’s rho between the number of structures present in an MSA against the maximum RMSD is 0.18, and against the mean RMSD it is 0.09. These very weak correlations show that in our dataset, the protein backbone conformation is well explored, with four or more sequence clusters with known structures. Therefore, we are not introducing a bias due to the available number of structures when considering the mean RMSD as a measure of the structural divergence of an MSA, this being the most robust measure to assess the structural divergence of a protein family.On the contrary, the correlation between the total number of structures and the mean change of the RASA is 0.54 and 0.49 against the fraction of changing contacts. Apparently, while the conformation of the backbone is well represented with four or more sequence clusters with known structures, this may not be enough to represent lateral chain changes between divergent structures. Such results show that the nature and location of lateral chains change greatly between structures, even maintaining the general fold (Russell and Barton, 1994). This is in agreement with Best and co-workers who concluded that more than 20–40 conformers are needed to explore the heterogeneity of side-chains (Best et al., 2006).

### 3.3. Correlation between residue solvent accessibility and conservation

An MSA can contain several proteins with available structures, but it is common practice to select only one structure from a selected reference sequence. As we have shown, protein structures in an MSA can significantly differ. For example, 50.63% of the MSA columns, on average, have residues exposed to the solvent in one structure but buried in another. It was found in previous works that the evolutionary rate of a site correlates with the residue RASA (Franzosa and Xia, 2009) and that conservation/variation scores (a proxy to evolutionary rate) negatively correlates with the solvent exposure of a residue (Echave et al., 2016; S.-W. Yeh et al., 2014), both conclusions were taken considering one structure per family. Knowing that the RASA changes considerably between structures, how does the correlation between the RASA and conservation/variation scores change considering different structures from the same MSA? We analysed the correlation between the RASA and the conservation (measured as KL divergence) of each structure in the MSA, as opposed to using only one structure for each MSA. We observed that the Spearman’s rho between the two variables varied over a wide range, from -0.82 to 0.23, with a mean of -0.37 (see distribution in Fig. 2A) and being -0.37 also the mean of the mean rho value per family. This means that for some structures, the RASA is strongly and negatively correlated with their family conservation while for others, it is not correlated or is even positively correlated. Indeed, the difference between the highest and the lowest correlation coefficient values (Δrho) within a family varies from 0.02 to 0.76, with a mean of 0.21 (Fig. 2B).This means that in some families, different structures show very similar correlations between the RASA and conservation (i.e. the difference is close to 0.02), whilst in other families, different structures may show very different correlation values (i.e the difference is close 0.74). The mean change of 0.21 can look small, but is proportionally important (57%) relative to the mean correlation value of -0.37.

**Fig. 2.**
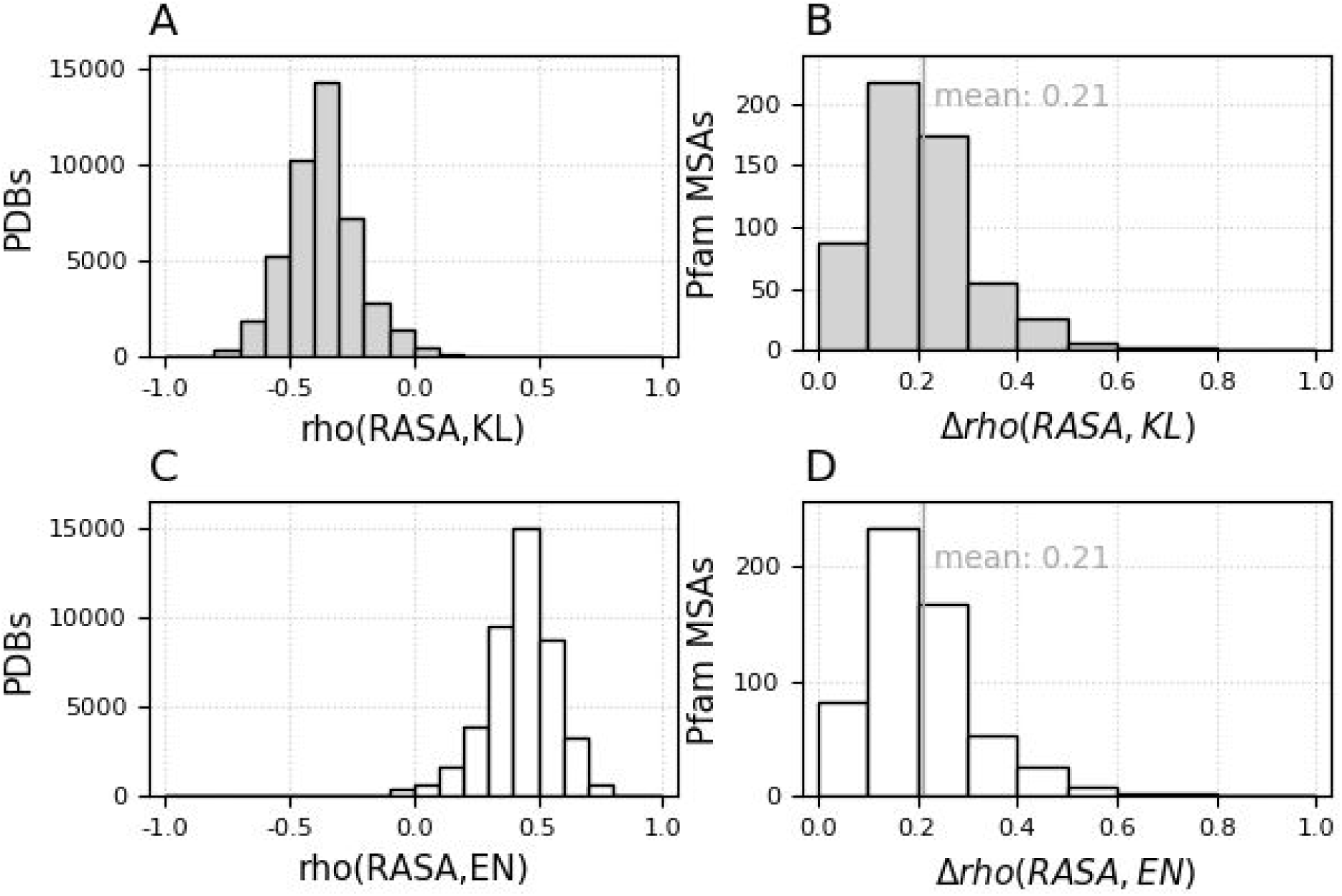
Spearman rank correlation coefficient between solvent accessibility (RASA) and conservation or variation. We use the Kullback-Leibler divergence (KL) as a measure of conservation (grey) or the Shannon entropy (EN) a measure of variation, inverse of conservation (white). **A and C)** For each PDB structure in our dataset we estimate the Spearman’s rho between the residue RASA and the conservation/variation measure of its MSA column. Histograms show the number of structures in our dataset per bin of rho. **B and D)** The same family has a value of rho for each available PDB. This histogram shows the number of Pfam MSA’s with a particular change (measuring the change as the difference between the highest and the lowest PDB Spearman’s rho within this family).

A previous study found a correlation value of 0.39 between the RASA and conservation, measured as 1 minus the Shannon entropy (EN) (S.-W. Yeh et al., 2014). In order to make comparable, we also calculated the correlation using EN as a measure of variation (instead of KL). The mean of the mean rho value per family between the residue RASA and EN is results 0.42 (the distribution is shown in Fig. 2C). Also, the difference between the highest and lowest correlation between EN and the RASA (Δrho) within a family varies from 0.02 to 0.78, with a mean of 0.21 (Fig. 2D), almost equal to results obtained using the KL divergence score (Fig. 2B).

### 3.4. Can more structures explain more of the conservation variance?

We then investigated whether the conservation pattern is better explained taking into account the solvent accessibility from several structures rather than from a single structure. To measure the solvent exposure of an MSA column, taking into account all the structures, we came up with two scores, the weighted mean RASA value and the weighted probability of a position to be exposed (see Materials and methods).

We used the Spearman rank correlation rho between KL and solvent exposure variables (since *a priori*, a linear correlation can not be assumed) using all families in the confirmatory dataset.The rho between the KL of each MSA column and the RASA of a single structure per family is -0.37 on average (see Fig. 2A), a little lower than the rho of -0.40 against the weighted probability of being exposed or the rho of -0.42 against the weighted mean RASA of the MSA column (Fig. 3). We found similar results using EN. In particular, the correlation values against the RASA values of a single structure, the mean RASA of a position or its probability of being exposed are 0.42, 0.48 and 0.46, respectively. The mean correlation values of EN or KL against the RASA of a single structure are significantly lower than the mean correlation values using information of multiple structures (*P* ⪡0.001 after a Wilcoxon signed rank test). That is to say,using the solvent accessibility from all available structures correlates better with conservation/variation scores than the RASA from a single structure.

**Fig. 3:**
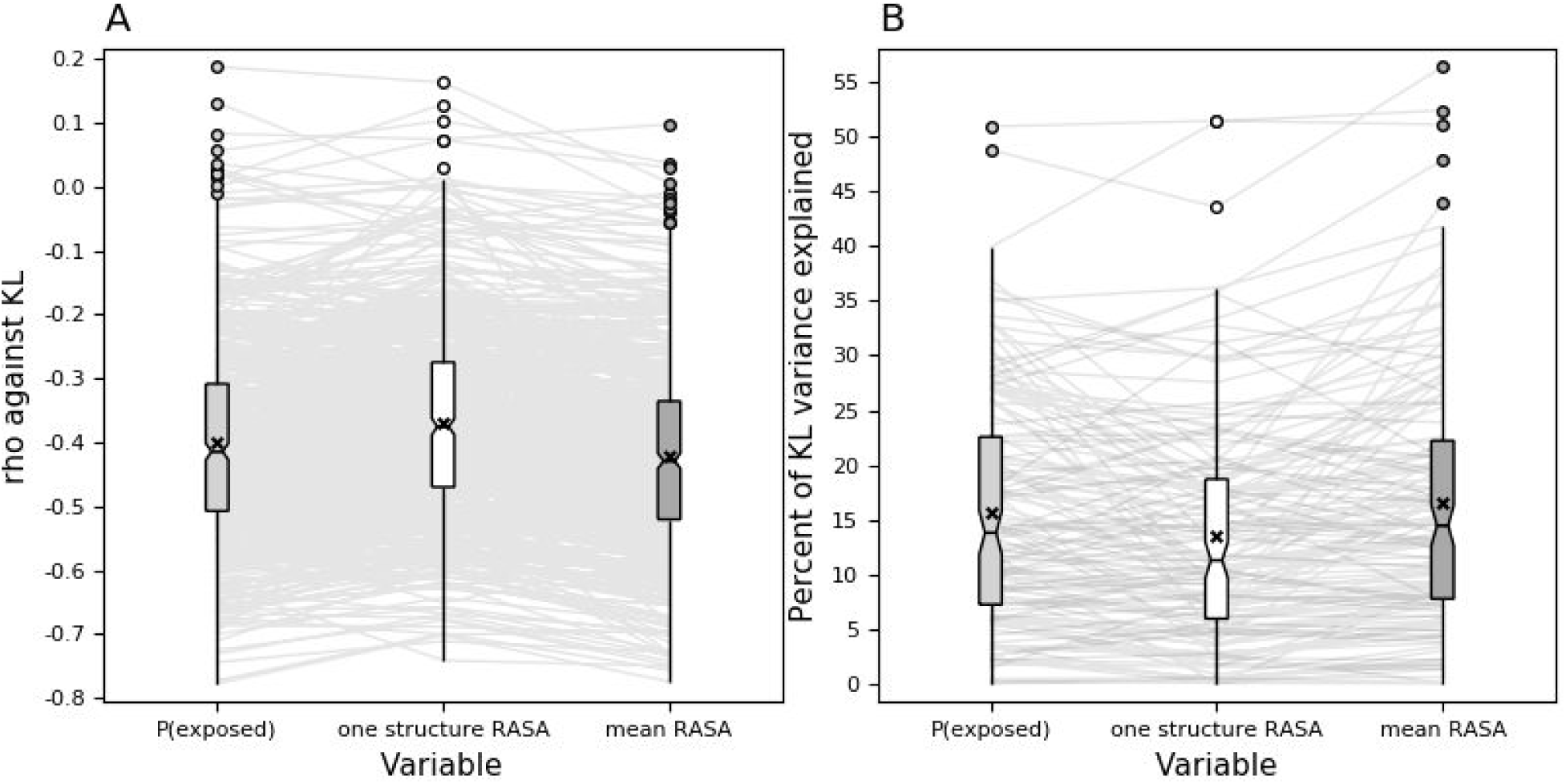
KL is better explained using the solvent accessibility of more than one structure. The boxplots show the distribution of the correlation scores between KL and P(exposed) and mean RASA. P(exposed) (light grey) and mean RASA (dark grey) include information from more than one structure. The distribution achieved using one structure from the MSA (white) is shown in the center. Each grey line connects the values of a protein family. A) The distribution of the Spearman rank correlation rho show more negative values when more than one structure is used. B) The percentage of KL variance explained is higher when more than one structure are used.

The relationship between conservation and the RASA, and related variables, is almost linear when the function *log(x+0.05)* is applied to transform KL values. We found that the RASA coming from a single structure for each MSA explains 13.47% of the variance, on average, (median 11.28%) using the squared value of the Pearson correlation coefficient, while the weighted probability of being exposed explains 15.57% of the variance, on average, (median 13.8%). The weighted mean RASA of each MSA column integrates data from many of structures and explains 16.53% of the variance, on average, (median 14.46%; Fig. 3 B). Therefore, using more than one structure, it is possible to explain between 2.1% to 3.06% more of the variance, on average, than using one structure from the MSA. Information is gained by including more structures, even taking into account the information lost due to mean and probability calculations. In fact, the fraction of KL variance explained using the weighted probabilities of being exposed is greater than using the RASA of a single structure for 63.09% of the Pfam families, and for 71.81% of the families using the weighted mean RASA values. Differences between the percentage of variance explained using either all the structures or a single representative show a low correlation with the number of available structures, Spearman’s rho 0.01 for P(exposed) and -0.06 for the mean RASA. Also, the same difference does not depend on the mean RMSD of the family, Spearman’s rho 0.15 against P(exposed) and 0.06 against the mean RASA. This demonstrates a lack of bias due to the number of structures, or structural divergence.

### 3.5. Covariation and protein contacts

It is a common practice to evaluate the performance of coevolution methods in terms of their ability to predict protein contacts; these usually have taken a single structure as a reference for a whole alignment (Buslje et al., 2009; Jones et al., 2012). But, how will the covariation scores perform for different structures when we have shown that ~53.32% of the inter-residue contacts change between proteins within a given protein family?

In order to address this question, we measured the Area Under the ROC Curve (AUC) to predict residue contacts for each structure in our dataset (with a PDB-Protein Data Bank-structure). The mean AUC for contact prediction was 0.74 for ZMIp and 0.76 for Gaussian Direct Coupling Analysis (DCA) (distributions are shown in Fig. 4A.). Surprisingly, the mean change in the AUC per family, defining change as the difference between the AUC with the PDB that performed highest and the one that performed the lowest, is only 0.029 for ZMIp and 0.031 for Gaussian DCA (shown in Fig. 4B.). Why is there almost no change in the predictive performance if almost half of the contacts change from one structure to the other?

**Fig. 4:**
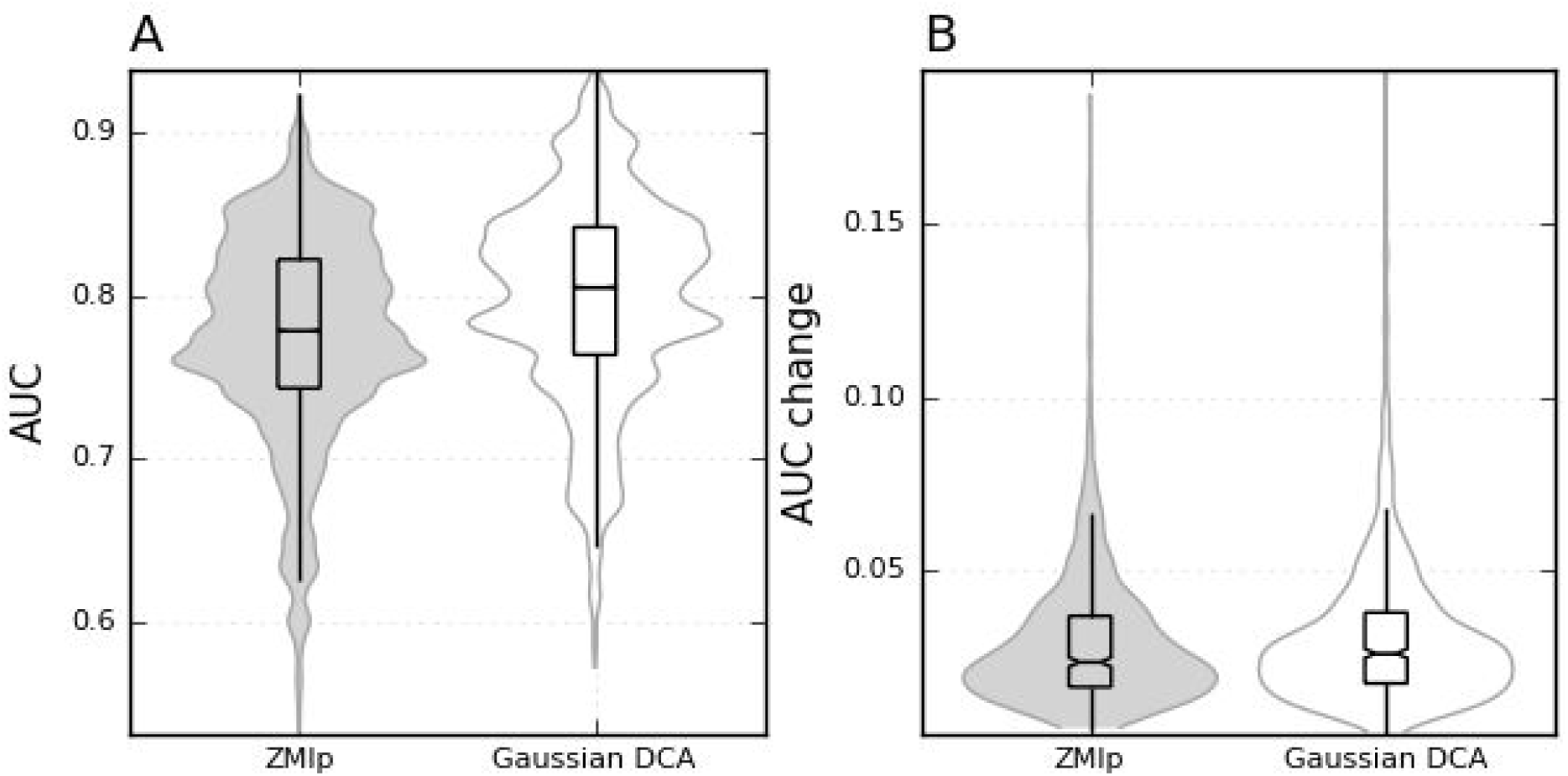
AUC for contact prediction by covariation methods. **A)** AUC distribution of the totality of protein contacts prediction for all PDB in our dataset. **B)** Distribution of the difference between the higher and the lowest AUC per family.

Subsequently, instead of using a contact matrix for each PDB, we defined contact matrices for each MSA. We considered that a pair of columns are in contact if positions are in physical contact in at least one structure; otherwise that column pair is considered not in contact. MSA columns in contact are then classified as conserved if they are present in all structures, and non-conserved or changing contact if the contact is absent in at least one structure.

The AUC to predict conserved contacts was 0.79 for ZMIp (Buslje et al., 2009; Zea et al., 2016) and 0.82 for Gaussian DCA (Baldassi et al., 2014). This indicates a good prediction performance for these methods in predicting conserved contacts through evolution. However, the AUC used to predict contacts specific for some structures (non-conserved) in the MSA is 0.66 for ZMIp and 0.67 for Gaussian DCA. This indicates a poor performance in the prediction of the conformer or subfamily specific residue contacts. Fig. 5 shows that the AUC to predict conserved contacts is always greater than the AUC to predict changing ones by ZMIp (Fig. 5A.) and Gaussian DCA (Fig. 5B.). In fact, for each Pfam family, on average, the conserved contacts are ranked higher than changing contacts or residue pairs not in contact as shown in Fig. 5.

**Fig. 5.**
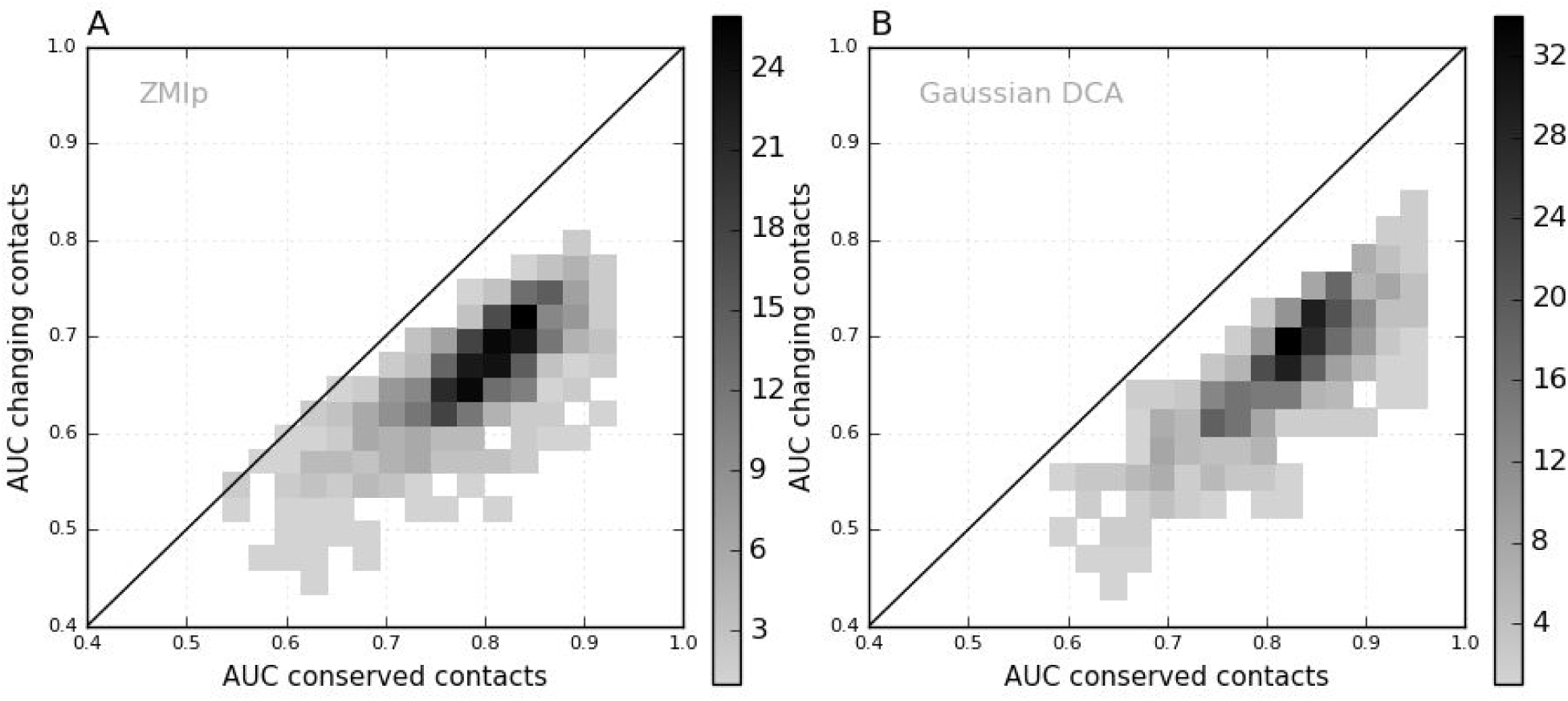
Covariation methods predict better the conserved contacts through the MSA. We calculated the AUC for conserved and changing contacts prediction for each Pfam family in our dataset. Each plot show a bidimensional histogram of the AUC to predict conserved contacts vs the AUC to predict changing contacts. Note that the performance to predict conserved contacts is DCA. higher since the cloud of points lies under the line y=x. **A)** Using ZMIp. **B)** Using Gaussian DCA.

This observation allows us to hypothesise that small changes in the AUC for different structures of a protein family can be caused by the ability of covariation scores to predict conserved rather than changing contacts. That is, the contacts that are common to all structures are the main contributors to the structural covariation signal, and therefore the ones that are predicted, equally, for all structures. If covariation scores were better in determining changing contacts than they presently are, values of the AUC would change much more after taking different structures.

### 3.6. Correlations of variables related to local packing density with conservation taking into account structural divergence

Next, we studied the relationships between conservation and structural variables when tested in large Pfam alignments containing multiple and diverse protein structures. We calculated the correlation score between conservation or variation of protein positions against structural information derived for each crystallographic structure in the family. To avoid problems derived from non-linear relationships, we used Spearman rank and partial correlation scores. In agreement with the pioneering work of Franzosa and Xia (Franzosa and Xia, 2009) and in opposition to more recent findings (S.-W. Yeh et al., 2014), we found that the RASA is the structural variable that shows the best correlation against conservation and variation scores. In particular, the mean Spearman 's rho between the RASA and KL is -0.37, while the correlation between the contact number (CN) or weighted contact number (WCN) and KL is 0.3 and 0.33, respectively. To avoid being misled due to data redundancy since different families contribute different numbers of structures, we sampled one structure per family 100 times and measured the average of the mean correlation values. The tendency did not change, neither for the correlation of KL against the RASA, nor for the CN and WCN. The average mean correlation values were -0.37, 0.31 and 0.34, respectively.

Results were similar using EN, a measure of variation, instead of KL. In particular, the mean correlation score between the RASA and EN was 0.43, while the mean rho between EN and the CN or WCN was -0.33 and -0.37, respectively. This tendency did not change when we sampled in order to avoid data redundancy, with the average correlation value out of 100 samples being 0.42, -0.33 and -0.37, respectively.

We found that in our dataset, the RASA correlated better with conservation or variation scores (KL and EN, respectively) than with variables related to local packing density (CN and WCN). In order to know if the correlation between the RASA and the evolutionary variables could be explained in terms of the local packing density scores, we measured partial correlations between the RASA and EN or KL by controlling for the CN or WCN. Results in Table 1 show that the correlation between the RASA and EN or KL did not disappear after controlling for the CN or WCN; they only show a small decrease (mean decrease was 0.15) in absolute value. This means that the contribution of the RASA to explain variation or conservation can not be completely explained in terms of local packing density scores.

**Table 1.**
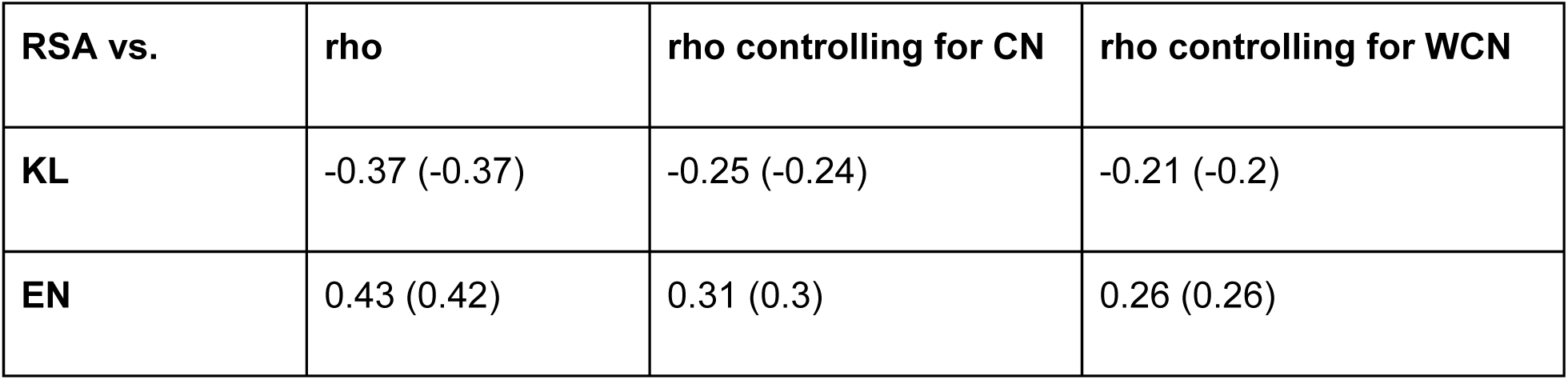
Spearman correlation scores between RASA and KL or EN, controlling by CN or WCN. The partial correlation score is calculated as *rho(RASA, row, column)*. So, the first column **RSA** simply the correlation coefficient without controlling for any variable. Numbers between parenthesis are the mean correlation scores after sampling 100 time only one structure per family in order to avoid bias by redundancy.

Previously, we showed that the mean RASA value across MSA columns showed the best correlation with the conservation of an MSA column (-0.42). In that sense, we tested if the mean CN and WCN also did better than a randomly selected structure from the MSA. We found a Spearman correlation coefficient of 0.32 between KL and the mean CN; this became 0.35 using a mean WCN instead of a mean CN. These two correlation values were only 0.01 unit, on average, above the average correlation values from sampling only one random structure for each MSA, 100 times. As can be seen in Table 1, the correlation between the RASA and KL of -0.37 was 0.06 units below the -0.43 correlation value between the mean RASA and KL. The same tendency in favour of the mean RASA can be seen using EN in place of KL divergence. In particular, the correlations between EN and the mean RASA, CN or WCN were 0.48, -0.34 and -0.38, respectively. We can conclude from these results that the RASA may be a better determinant of residue conservation than local packing density scores.

Other research groups highlight the importance of local flexibility as a determinant of evolution rates (Huang et al., 2014). We compared the contribution of structural flexibility, taking into account all structural information available for the MSA, against the contribution of the RASA using correlation coefficients. In particular, we compared our correlation between KL or EN against the mean RASA per MSA column (which were -0.42 and 0.48, respectively), against the correlation scores of KL or EN against the Root Mean Square Fluctuation (RMSF) of an MSA column, which were -0.33 and 0.34, respectively. This last correlation value of 0.34 decreased to 0.16 when the mean RASA was taken into account (partial correlation). Because the correlations of the evolutionary scores are lower in absolute value against RMSF than against the mean RASA, and because almost a half of the correlation value against RMSF can be explained by the mean RASA, we can conclude that solvent exposure appears to be a better determinant of conservation than flexibility.

### 3.7. Case example: Structural change of a protein kinase domain

The serine/threonine-protein kinase PIM-1 (UniProt: PIM1_HUMAN) is a proto-oncogene involved in cell survival and proliferation (Tursynbay et al., 2016). Its biological activity is determined by its protein kinase domain (Pfam: PF00069). We analysed 615 structures (only the columns with solved residues in all structures). To avoid redundancy, the structures were clustered at 0.4 Å of RMSD giving 335 clusters.

The protein kinase domain Pfam alignment has 179123 sequences that make up 37287 clusters at 62% identity. Only 48 sequence clusters have structural information and the 615 structures are contributed by 77 proteins. After removing insert columns and columns without structural information, our alignment ended up with 165 columns. The maximum RMSD between the most dissimilar structures (maximum family structural divergence) was 12.05 Å, achieved between two protein domains with 34.55% sequence identity (see Fig. S4.).

In our analysis, we had 80 conformers available for PIM-1, showing a maximum RMSD between two conformers of 1.31 Å. Nevertheless, another family member, the human Aurora kinase A,had a maximum RMSD between two conformers of 8.37 Å (see Fig. S4.).

The structural alignment implicit in the Pfam sequence alignment is shown in Fig. 6. The mean RMSD between the structures in our dataset was 3.39 Å. This structural divergence in the C alpha backbone was accompanied by large changes on the residue’s lateral chain position. We found that 85.45% of the MSA columns have residues that change their buried or exposed in at least one structure. Fig. S5. shows how the PIM-1 4N70 structure residue RASA are related to the mean RASA of each column of the MSA. We also found that 81.26% of protein contacts (where 100% is the number of contacts that exist in at least one structure), disappeared in at least one structure of the MSA. Fig. S6. shows the top 5% Gaussian DCA scores and the probability of each pair of positions being in contact through the alignment. The AUC to predict conserved contacts (pairs with a 1.0 probability of being in contact) by Gaussian DCA score in this particular family was 0.924, while the score to predict contacts that change through the alignment was 0.742. Fig. S7. shows the probability of a position pair being in contact as a function of the Gaussian DCA score. Notably, the top 1% Gaussian DCA scores pointed out position pairs with a high probability of being in contact.

**Fig. 6.**
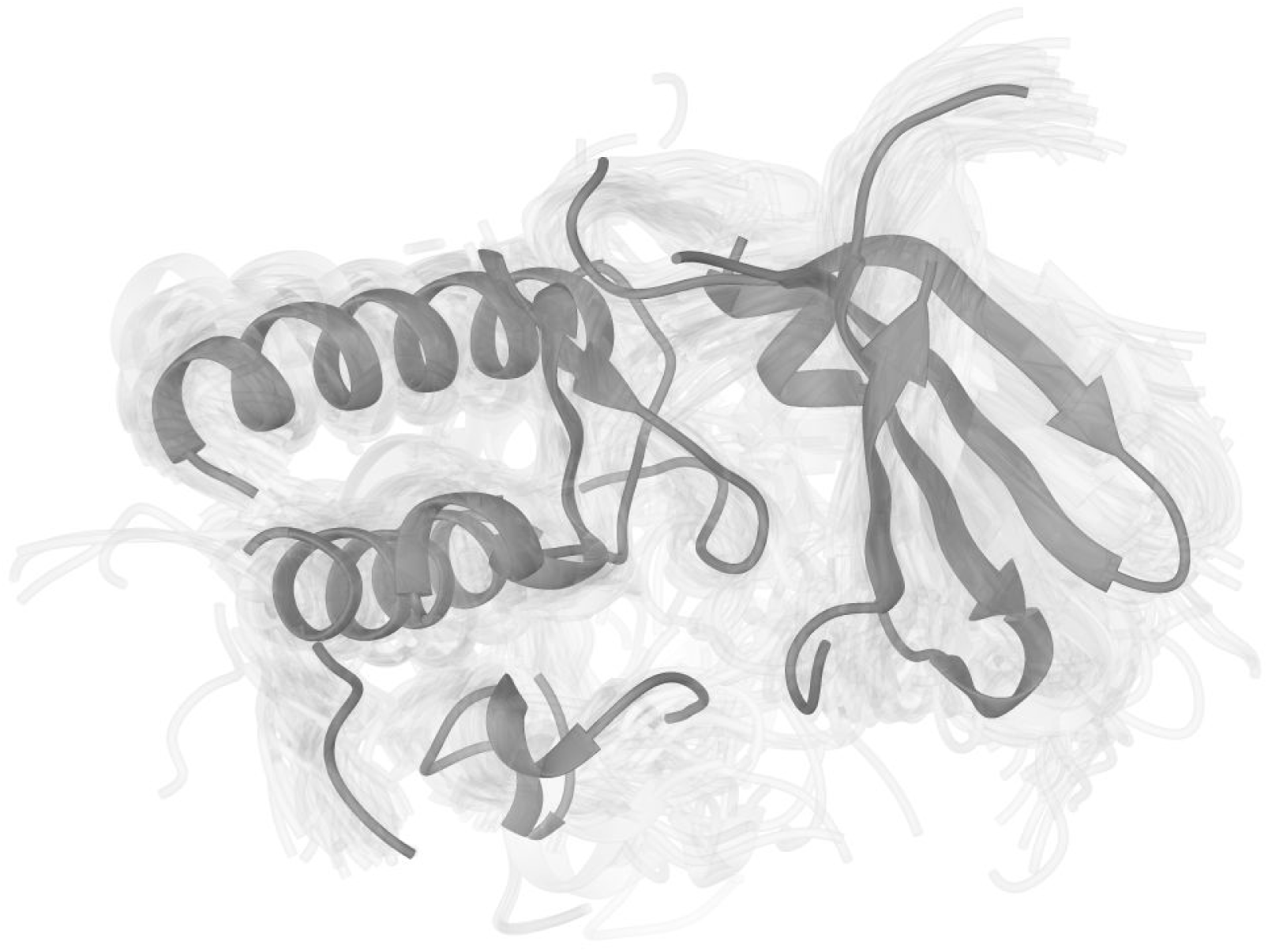
Structural superimposition of protein kinase domains. One PDB for each of the 77 protein domains with known structures in the protein kinase family was selected for this visualization. The structure in dark grey corresponds to the human PIM1 (PDB 4N70). The other structures are shown in light grey.

## 4. Discussion

It is common a practice to map sequence information derived from homologous proteins into one of the structures of a protein family. Sometimes a known conformer from the native state of the studied protein is used, and, at other times, a conformer from a close homologous protein. However, a number of potential problems occur that lead to the following questions: Is the native state of the studied protein fully represented by a single conformer structure? Are close homologous native states similar enough to use one instead of the other? When the sequence information to be mapped comes from a multiple alignment of homologous sequences, further questions arise:

1. Can a single conformer from a single structure represent the structural space of the protein family?
2. Is the native state of each protein in the alignment contributing the same structural constraints during protein evolution?
3. Can a single structure be enough to represent the structural constraints modulating the substitution pattern in the different positions of an whole protein family?

In this study, we have shed some light on the latter questions. In particular, we studied how structural divergence is related to evolutionary information.

In particular, we analysed two types of evolutionary information that could be extracted from a multiple alignment of homologous sequences: conservation and coevolution. Conservation can be seen as evidence of evolutionary constraints on single positions, while coevolution refers to affecting two or more positions at the same time. A large number of distant homologous are needed for some of these methods, particularly covariation methods used to estimate coevolution. Consequently, Pfam alignments are a common choice for coevolutionary studies given their high quality and the large number of distantly related homologous sequences. We found that Pfam alignments also showed a large structural divergence. We understand structural divergence as being a series of structural changes due to sequence divergence through the accumulation of nonsynonymous substitutions. Even when sequences inside a Pfam family share a structural fold, large RMSD values are obtained as derived from backbone comparisons. Also,subtler structural features, for example residue contacts and solvent accessibility, change even more between structures derived from the same multiple sequence alignment. These changes were evident in the case example referring to the protein kinase domain, where structures show the same fold; however, many structural changes are observed, even at the conformational diversity level. Since Pfam families show large structural divergence, a single protein structure can not represent all the structural space of its family.

Structural constraints to the evolutionary process exist in order to maintain the native state of proteins (Zea et al., 2013). In particular, residue solvent accessibility has been declared a structural determinant of the evolutionary rate. In short, residues on the surface tend to vary more than residues in the core of the structure (Franzosa and Xia, 2009). Also, residue contact number has been correlated with residue conservation (S. W. Yeh et al., 2014). In addition, the need to maintain inter-residue contacts imposes constraints to position pairs variation, leaving a covariation signal (Dunn et al., 2008). What is more, conservation and covariation signals are recovered from multiple sequence alignments with a large structural space. How does this structural space constraint protein evolution?

To understand this, we measured how the correlation between solvent accessibility and constraints conservation, and the predictive performance on contact prediction of covariation scores, changes when different structures from the same family were used. Different outcomes were expected because of the large structural space of protein families. We found that the correlation between solvent accessibility and conservation/variation scores greatly changed depending on the selected structure, as expected. However, the predictive performance of covariation scores to predict contacts almost did not change between structures.

For each multiple sequence alignment, we have a vector of conservation/variation values, with a value per position. A conservation vector is unique for a given protein family. However, for each structure in the family, we have a RASA vector with a RASA value per residue in the structure. A correlation value can be measured between the conservation vector and each RASA vector. Previous studies used a single structure to explore this structural feature, so they had a single correlation coefficient per family. The approach of these previous studies is correct and can be thought of as stratified sampling, where a single random structure is selected from each stratum (or protein family in this case). In fact, reported mean correlation values in these datasets are very similar to mean correlation values in our dataset using all available structures.However, an interesting fact we have shown, is that correlation values between solvent accessibility and conservation can vary greatly inside a protein family. The magnitude of the change between correlation values in a family is directly related to the magnitude of the change in solvent accessibility values of its known structures. Solvent accessibility changes considerably in the structural space of a protein family. In fact, positions that are buried in one structure of the family can be totally exposed in another structure. If solvent accessibility determines conservation, we may expect that a measure taking into account all known structures can give a better picture of the structural constraint. Indeed, we found that measures which resumed the solvent accessibility of all known structures were better at explaining the conservation calculated the multiple sequence alignment.

To capture the solvent accessibility diversity in a protein family in a single vector, we calculated two measures: the probability of a residue of being exposed to the solvent and the mean RASA value of the position. Using these parameters, we found that the conservation variance explained, was 16.5% on average, slightly above the value found when only one structure was used per family (13.5%). The small increment in the explained conservation variance may be due to the fact that structural factors are not the main constraints to sequence divergence during evolution (Bloom et al., 2006), that almost half of the columns do not change their exposed/buried status, on average, or that the structural space of these protein families are still incomplete.

Additionally, we also analysed other structural variables that correlate with conservation scores, in particular, variables related to contact density such as contact number and weighted contact number. These variables show lower correlation values than solvent accessibility, even when multiple known structures are used. A gain in correlation values between conservation and contact density does not occur when more than one structure is used. That is to say, the correlation does not increase when the mean value per column of these variables is used instead of the value from a single structure.

Other variable derived from the known structural space of a protein family is the Root Mean Square Fluctuation of the alpha carbons (RMSF). This variable, related to protein flexibility through evolution, shows lower correlation values against conservation scores than the mean RASA of alignment columns. Therefore, contact density and flexibility variables may be poorer determinants of residue conservation than solvent accessibility in our dataset.

It is known that residue contacts can lead to a covariation signal, i.e. coevolution (Dunn et al.,2008). In fact, the predictive performance measure of the covariation scores is based on this from idea. We observed that many inter-residue contacts changed through evolution and, consequently, we expected predictive performance changes when different structures were used. However, our results showed otherwise. The fact that the predictive performance of covariation scores remains the same between structures can be explained by the fact that these scores predict conserved contacts, i.e. inter residue contacts in common. This result is in concordance with recent findings (Rodriguez-Rivas et al., 2016) where evolutionary conserved contacts between proteins were predicted using covariation methods.

As a result of a covariation measure ranking first conserved contacts, it is difficult to predict conformer or subfamily specific contacts (i.e. functional important contacts that change between structures). For covariation scores, the common practice of using a single structure to measure predictive performance is not misleading. However, consideration of the large structural space implicit in diverse multiple sequence alignments can be useful in the development of new contact prediction methods more oriented to the prediction of contacts that change between conformers or through evolution.

In this study, we used a large dataset of Pfam alignments. These multiple sequence alignments are commonly used to measure covariation scores because they contain a large number of remote homologous. The large number of evolutionarily diverse homologous sequences with known and accurate structural information also make these alignments a good dataset in which to explore our questions. Although such Pfam alignments may represent the more extreme case of structural divergence, large structural differences can be seen at high sequence identity. Infact, residue contacts and solvent accessibility can greatly change, even between conformers of the same protein. Because of this, we believe that evolutionary information derived from protein alignments, even for smaller and curated alignments, deserve to be matched with all known structural information available in order to achieve a better understanding of the studied system, when a single protein or protein family is analysed.

When no or little structural information is available, we must bear in mind that conservation and covariation measures take into account the implicit structural space of the sequences used to calculate these scores. In particular, residue conservation is not directly related to the RASA of a single protein structure. Therefore, mapping conservation scores to the surface of a single structure could potentially lead to misinterpretations. In the case of covariation scores, high values tend to indicate conserved residue contacts in the family. Therefore, interpreting these values as protein/conformer specific may be incorrect.

In summary, when analysing a single protein family, we can be more confident in the information provided by all its available structures rather than from a single representative structure. The weighted mean RASA of an MSA column after a structural clustering, the weighted probability of a residue being exposed through the alignment and the probability of a pair of residues being in contact give good insights into understanding conservation and coevolution (or at least covariation) patterns in the MSA.

## 5. Conclusions

It is a common practice to correlate evolutionary information derived from protein family alignments with a single structure taken as reference. We have shown that matching evolutionary information with all the available structural information provides a better understanding of the studied system. This is particularly important when a single protein or protein family is analysed.

We found large structural changes between proteins within a Pfam family at the residue level. In particular, residue solvent exposed area and inter residue contacts change greatly between within a protein family. As a result, linking evolutionary information as conservation and coevolution to variables calculated from a single structure, can be misleading. The use of solvent accessibility data from multiple structures permits a better understanding of the conservation pattern than a single structure. Also, solvent accessibility is a better determinant of residue conservation when multiple structures are taken into account, compared to flexibility or residue contacts.

Covariation methods, a proxy to coevolution, are less sensitive to differences in the residue contacts of different protein structures. This is because they predict with fitter performance structural contacts that are conserved through the alignment than protein specific contacts. In general, our results show that the use of multiple structures from a protein family enriches the capture of the structural constraints affecting protein evolution.

## Acknowledgements

We would like to thank the members of the Structural Bioinformatics Unit of the Fundación Instituto Leloir and members of the Structural Bioinformatics Group of the Universidad Nacional de Quilmes for their multiple comments and useful discussions. All the authors are researcher of the Argentinean National Research Council (CONICET). This work was supported by Universidad Nacional de Quilmes [grant UNQ 1402/15] and CONICET [grant PIP 1087].

